# Ovipositor morphology and mechanosensory innovation traits favor niche width expansion in *Drosophila*

**DOI:** 10.64898/2026.04.29.721748

**Authors:** Shan He, Tianpeng Wang, Yuan-jun Yu, Xiao-nan Yang, Wu-fan Zhang, Bin-yan Lu, Yi-bo Luo

## Abstract

Ecological niche breadth is key factor associated with adaptive radiation and speciation. For most *Drosophila* species, the firm surface of intact ripe fruit acts as a physical barrier to oviposition, effectively restricting them to the saprophagous resources. How species overcome such mechanical constraints at the behavioral and sensory levels, and whether doing so leads to niche breadth expansion or niche specialization, remain poorly understood. Here, using comparative behavioral assays across ten *Drosophila* species, we show that substrate physical hardness is a critical barrier preventing most species from exploiting ripe fruit. Among species capable of puncture oviposition, there are two distinct evolutionary strategies among *Drosophila* species. With its serrated ovipositor and strong preference for fresh fruit, *D. suzukii* represents a case of ripe-fruit specialization. In contrast, *D. immigrans* with the needle-like ovipositor, gains access to fresh fruit while retaining the ability to exploit decaying substrates, thereby expanding its resource niche. Thus, *D. suzukii* considered as the typical evolutionary niche specialization, while *D. immigrans* represents niche expansion. Furthermore, using the genetic toolkit of *D. melanogaster*, we further identify the *Inactive* (*IAV*) mechanosensory channel as important regulator of stiffness-dependent oviposition inhibition. Loss of *iav* reduced inhibition on firm substrates, while cross-species rescue experiments showed that *IAV* orthologs differed in their ability to restore this response in a common genetic background. Puncture oviposition was also associated with successful offspring development on firm fruit and with the use of firm-surfaced hosts under natural conditions. Together, these findings identified a novel niche expansion of *D. immigrans* among the saprophagous *Drosophila*.

## INTRODUCTION

The evolution of ecological niche breadth is central to adaptive radiation and speciation (Nyman, 2010; Sexton et al., 2017). When adapting to novel environments, herbivorous insects generally follow one of two distinct evolutionary trajectories: niche specialization or niche expansion. Niche specialization typically involves an evolutionary trade-off, functioning as a niche shift where a lineage gains the ability to exploit a novel resource but concurrently loses the capacity or preference for its ancestral hosts (Forister et al., 2015; Sexton et al., 2017; Sjödin et al., 2018). In contrast, niche expansion occurs when a lineage successfully incorporates a novel ecological space while retaining its ancestral resource base (Futuyma and Moreno, 1988; Sjödin et al., 2018). In either case, successful niche exploration requires complex coordinated changes in behavior, physiology, and sensory processing (Auer et al., 2020; Riffell, 2020). Understanding these transitions matters for both evolutionary ecology and agriculture, because adaptation to crop and food resources can directly promote pest emergence.

For fruit-breeding insects, the fruit-ripening continuum defines a major resource axis along which niche breadth varies. As fruits progress from fresh to rotten, they change in volatile composition, sugar content, and organic acid balance, but also in surface texture and penetrability (Basson et al., 2010). Although most work on host shifts has focused on the chemical dimension of this gradient (Auer et al., 2020; Dweck et al., 2021; Karageorgi et al., 2017), the physical dimension of the ripening gradient represents an independent ecological axis that may impose a different kind of constraint. Ecological theory distinguishes between substitutable and non-substitutable resources, showing that competition and niche structure differ fundamentally depending on whether one resource axis can compensate for another (Ashby et al., 2017). In this framework, chemical features of fruit are relatively substitutable for oviposition-site choice, because insects can evolve shifts in sensory tuning to track different volatile profiles while still locating acceptable substrates. By contrast, the intact skin of fresh fruit acts as a non-substitutable barrier: female insects may detect and even be attracted to fresh fruit chemical cues, yet still be unable to use that resource if they cannot physically penetrate the oviposition site (Atallah et al., 2014; Keesey et al., 2015). How insects overcome this mechanical gate, and at what behavioral, morphological, and sensory levels this occurs, remains poorly understood.

The genus *Drosophila* provides a comparative framework for studying host and substrate niche adaptation. Closely related species often breed on distinct substrates, and these behavioral differences can be tested using controlled oviposition assays, supported by the genetic tools for mechanistic analysis in several model species (Anholt et al., 2020; Durkin et al., 2021; Markow and O’Grady, 2005). Most drosophilids are saprophagous, relying on decaying soft-surfaced fruits for oviposition and larval development (Markow and O’grady, 2008). Previous work on *Drosophila* niche exploration has focused on chemical barriers, with well-studied cases including *D. sechellia* and *D. yakuba*, which have evolved as niche specialists on *Morina citrifolia* fruit (Auer et al., 2021; Erlenbach et al., 2023; Louis, 1986; Rkha, 1991; Yassin et al., 2016). Physical barriers to oviposition, however, have received less attention as drivers of niche expansion. Within the subgenus *Sophophora, D. suzukii* represents the best-known exception to the saprophagous preference: it has evolved an enlarged, serrated ovipositor that allows females to pierce intact fruit skin, and it oviposits preferentially on fresh rather than rotten fruits (Atallah et al., 2014; Green et al., 2019; Karageorgi et al., 2017). *D. suzukii* shows strong preference for ripe fruit and reduced responses to late fermentation-associated substrates, representing this transition as a niche specialization trajectory (Karageorgi et al., 2017). By contrast, within the subgenus *Drosophila*, *D. immigrans* has been reported to breed on firm-skinned fruits such as citrus and strawberry, where it can also cause agricultural damage (Atkinson, 1981; Živković et al., 2019). Importantly, *D. immigrans* also breeds on rotting fruits, vegetables, and mushrooms at multiple stages of decay, suggesting that it retains broad ancestral resource use while gaining access to fresh fruit (Atkinson and Shorrocks, 1977; Patterson and Stone, 1952). Thus, although *D. suzukii* and *D. immigrans* both oviposit on firm fresh fruit, they appear to differ fundamentally in niche breadth: *D. suzukii* is a ripe-fruit specialist, whereas *D. immigrans* may represent true niche expansion across the ripening gradient.

Oviposition site selection in *Drosophila* is a multisensory decision-making process in which females integrate olfactory, gustatory, visual, and mechanosensory signals before laying eggs (Yang et al., 2008). Changes in sensory receptor genes have been repeatedly linked to host shifts across drosophilid systems. For example, modifications to olfactory sensory receptor genes are associated with the transition to herbivory in *Scaptomyza flava* (Goldman-Huertas et al., 2015), specialization on morinda fruit in *D. sechellia* (Auer et al., 2020), and ripe-fruit use in *D. suzukii* (Karageorgi et al., 2017; Xue et al., 2025). Similarly, changes in gustatory receptor genes can also shape oviposition decisions, as seen in the altered bitter and sugar receptor genes associated with host use in *D. sechellia* and *D. melanogaster*. (Reisenman et al., 2023). By comparison, the contribution of mechanosensory systems to host shifts remains less well understood. In *Drosophila*, mechanosensory neurons are distributed across the labellum, tarsal leg segments, ovipositor bristles, and internal reproductive tract (Gou et al., 2014; Hehlert et al., 2021), and several ion channels expressed in these neurons contribute to stiffness discrimination (Hehlert et al., 2021; Wu et al., 2019; Zhang et al., 2016). Genomic and transcriptomic studies have suggested that mechanosensory genes have changed in *D. suzukii* relative to generalist species, but the functional consequences of those changes remain unclear (Durkin et al., 2021). How mechanosensory signals input regulates oviposition decisions on stiff substrates remains poorly understood. Moreover, it is unclear how reduced behavioral inhibition interacts through special morphological ovipositor of *D. immigrans* to facilitate this species use intact fresh fruit while retaining its broader ancestral resource range.

In this study, we investigate how *D. immigrans* incorporates intact fresh fruit into a broad resource niche. We have combined behavioral assays, comparative morphology, functional genetics, and field surveys across ten *Drosophila* species to dissect the mechanisms of ripe-fruit niche expansion. First, we test how substrate hardness shapes oviposition decisions across species while decoupling stiffness from chemical cues, allowing us to identify the behavioral basis of puncture oviposition. Second, we examine the morphological and molecular basis of stiffness sensing, using the genetic toolkit of *D. melanogaster* to identify essential mechanosensory channels, with particular focus on the *Inactive* (*IAV*) channel, and to test whether functional divergence in this gene contributes to species-specific behavior. Third, we ask whether overcoming the mechanical barrier of fresh fruit confers a direct fitness advantage and predicts ecological host use under natural conditions. By comparing these species within a broader drosophilid framework, we reveal how mechanical innovation can expand ecological niche breadth while preserving ancestral resource base.

## RESULTS

### Oviposition preferences and behavior to different fruits substrates across *Drosophila* species

To determine the oviposition preference of ten different fly species between ripe fruits and rotten fruits, we used a two-choice oviposition assay to compare the oviposition preferences of flies between intact ripe and rotting strawberries (Figure 1A). These species exhibited divergent oviposition behavioral preferences when offered fresh and rotten fruit substrates. Seven of ten species, including *D. melanogaster* and its close relatives, exhibited a strong and significant preference for the rotten substrate (Figure 1A). On the contrary, the specialized *D. suzukii* displayed an opposite strong and significant preference for fresh strawberries, consistent with its known status as an invasive pest of fresh fruit. Surprisingly, both *D. immigrans* and *D. biarmipes* showed no significant oviposition preference for either fresh or rotten fruits. The oviposition preference of *D. immigrans* was highly consistent across individuals, with most data points clustered preference index near to zero, indicating that flies treated both fresh and rotten fruits as equally acceptable. However, *D. biarmipes* displayed high individual trials variability ranging from a strong preference for rotten fruit to a preference for ripe fruit (Figure 1A). These results showed that *D. immigrans* and *D. biarmipes* act as behavioral generalists, unlike the specialist *D. suzukii*, they accept both intact ripe and rotten fruits for oviposition.

**Figure 1.**
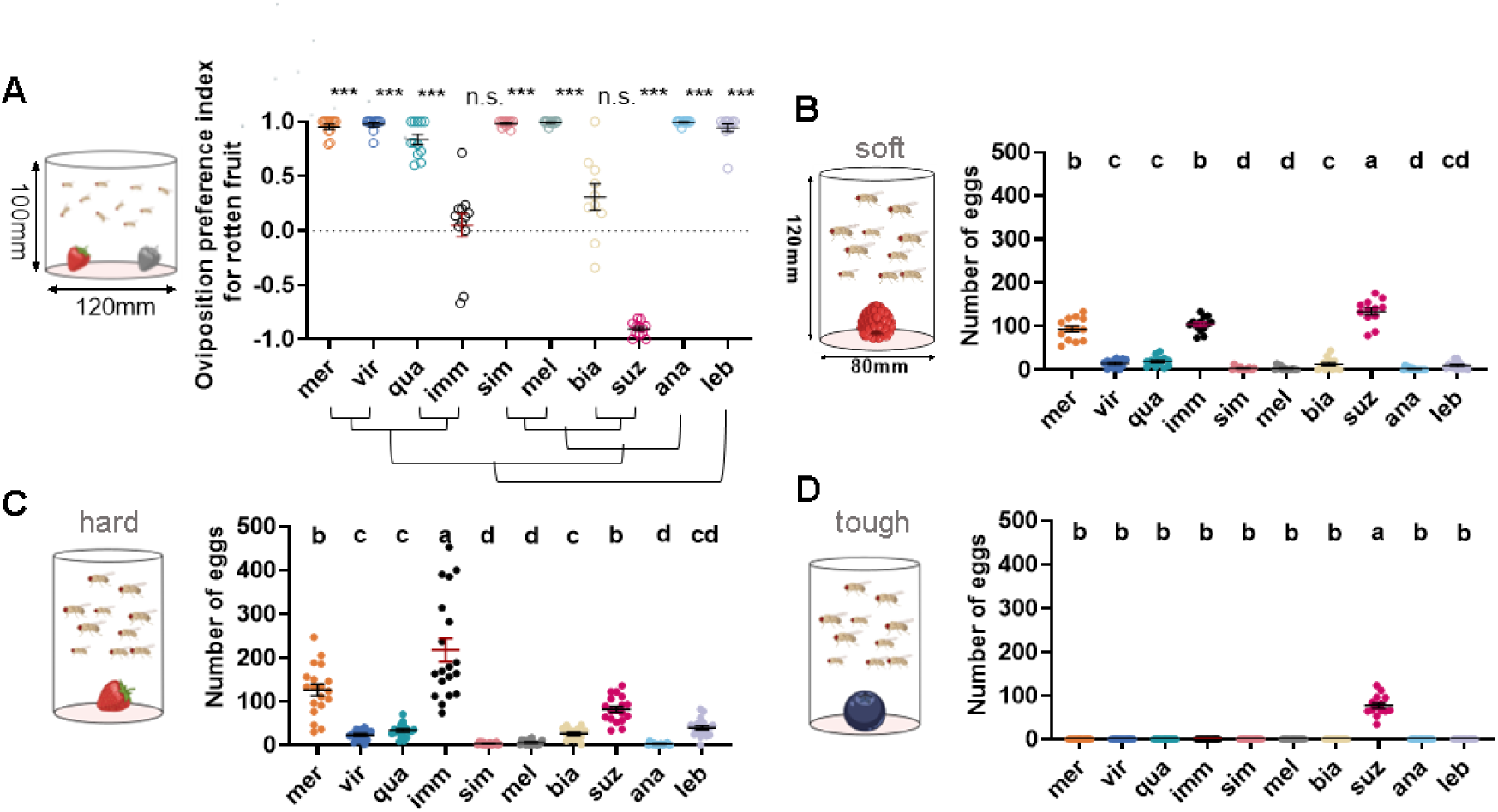
Oviposition behaviors on substrates with different stages of fruit maturation differ among 10 species. (A) Oviposition preference index (OPI) of ten fly species in two-choice assay between ripe and rotten fruits (strawberry). Schematic of two-choice oviposition preference assay is shown on the left. D. suzukii prefers to oviposit in ripe fruit, *D. immigrans* and *D. biarmipes* show no significant preference for either fruit, and other 7 species (e.g. D. melanogaster) prefers rotten fruit. Preference P-values were calculated using a Wilcoxon signed rank test against a theoretical value of 0 (no preference). mer-D. mercatorum, vir-*D. virilis*, qua-*D. quadrilinate*, imm-*D. immigrans*, sim-*D. simulans*, mel-*D. melanogaster*, bia-*D. biarimpes*, suz-*D.suzukii*, ana-*D. ananases*, leb-*S. lebanenonsis*. Error bars are SEM. n=12. ***, p≤0.001; **, p≤=0.01; *, p≤0.05; ns, p>0.05 for all future figures. (B) Mean number of eggs laid by ten fly species in a no-choice assay on single fruits, representing a range of surface stiffness (B: raspberry-soft skin, C: strawberry-hard skin and D: blueberry-tough skin). Most species exhibit a significant reduction in egg-laying on harder fruits, whereas *D. suzukii* and *D. immigrans* maintain high oviposition rates. Each data point represents one experimental trial. Significant differences are denoted by letters (P<0.05), treatments sharing a letter are not significantly different. Two–way ANOVA followed by Tukey’s multiple comparisons test. Error bars are SEM. n=12 – 14 replicates per condition.

Then, we used a no-choice assay to quantify the number of eggs laid by ten flies when provided with only one type of fruit, representing a range of surface stiffness: raspberries (soft skin), strawberries (hard skin), and blueberries (tough skin) (Figure 1B-D). When only fresh fruit was provided, most species significantly reduced their egg-laying rate (defined as the mean number of eggs laid by females within a 24-hour window) (Figure 1B-D). Again, *D. suzukii* exhibited a high oviposition rate across all three fresh fruit types. *D. biarmipes* only laid a few eggs on raspberries, which have relatively soft skin, but showed little activity on both strawberries and blueberries with harder skins. Among the other species, *D. immigrans* and *D. mercatorum* exhibited high oviposition rates on raspberries. However, on strawberries with hard skin, *D. immigrans* laid the highest number of eggs, surpassing even *D. suzukii* (Figure 1C). Curiously, the oviposition rate of both *D. immigrans* and *D. mercatorum* dropped significantly on the blueberries which having tough skin, the highest surface stiffness among three kind fruits (Figure 1D).

Considering the complex chemical signals of different fruits which could influence oviposition behaviors of *Drosophila* species, we developed the agarose gels with varying concentrations as the oviposition system. We first quantified the stiffness of fruits surface from ripe to decayed stages (Figure 2A). We found that fresh fruit surface of blueberries had harder surface than that of raspberries and strawberries. As these fruits decayed, their surface stiffness decreased and reached a low stiffness level (Figure 2A). We measured the stiffness of decaying fruit surface is about 0.25%-0.5% agarose, while the fresh fruit surface of blueberries is about 1.0%–2.0% agarose (Figure 2A).

**Figure 2.**
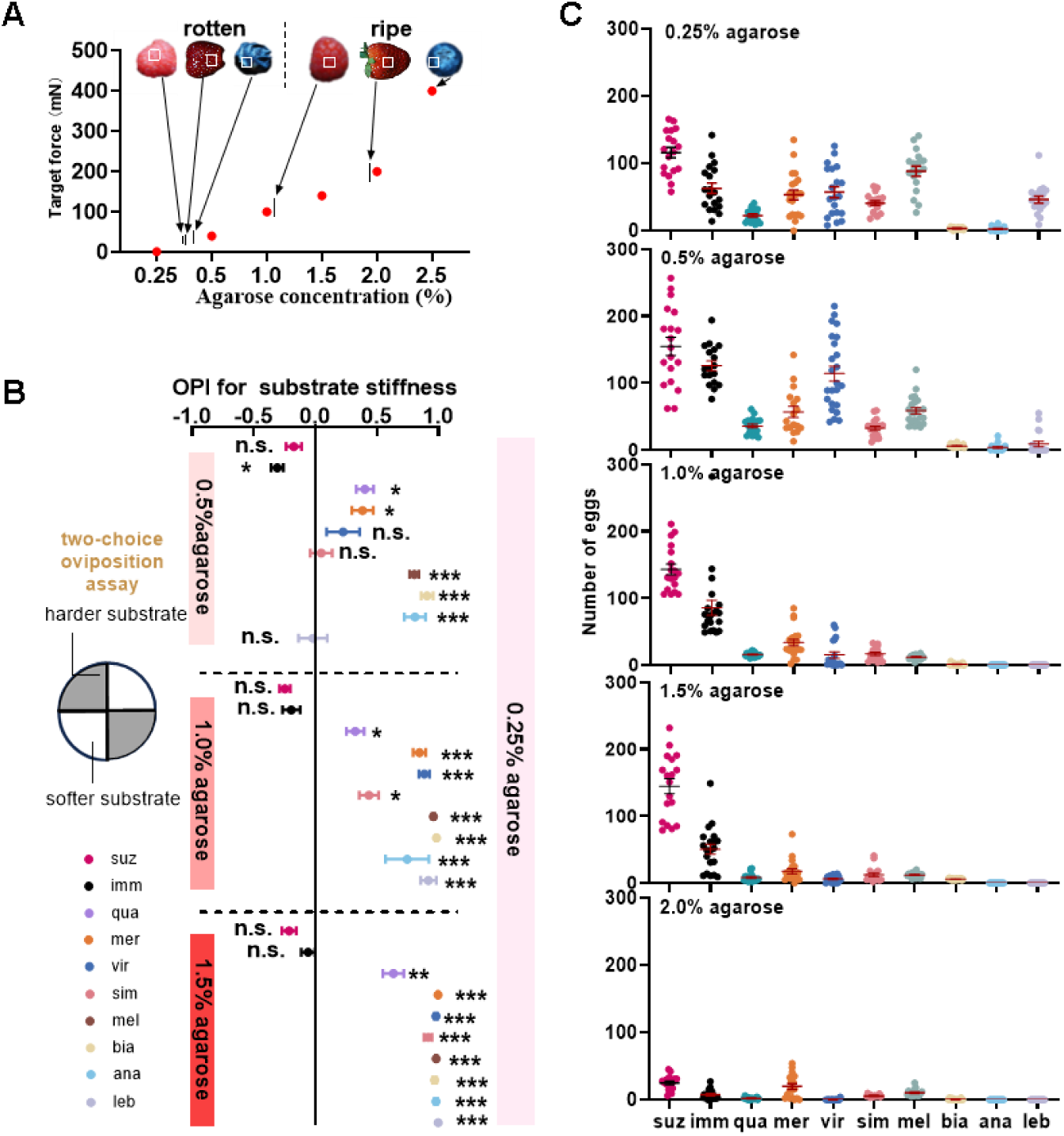
*D. immigrans* shows stronger preference and tolerance for stiff substrate or substrate stiffness. (A) Graph of substrate stiffness of substrates of different agarose concentrations (red dots) and fruits (gray arrow). (B) Oviposition preference index for 0.25% agarose of ten fly species at difference agarose ranges (0.5%, 1.0% and 1.5% agarose). While most species show a strong aversion to harder substrates, *D. immigrans* and *D. suzukii* show no significant preference or slight preference for harder agarose. P values were calculated via Dunnett’s multiple comparisons test against a theoretical value of 0.25%. Error bars are SEM. n=16-20. (C) The numbers of eggs laid by ten fly species in a no-choice egg-laying paradigm on substrates of the indicated agarose concentrations (0.25%, 0.5%, 1.0%, 1.5%, and 2.0% agarose). *D. suzukii* and *D. immigrans* maintain high egg-laying rates even at hard substrates (1.0% and 1.5% agarose), whereas other species showed oviposition rates drop sharply. Two-way ANOVA followed by Tukey’s multiple comparison test. Values indicated with different letters are significantly different (p<0.05). Error bars are SEM. n= 18-22.

We then selected four agarose concentrations (0.25%, 0.5%, 1.0%, and 1.5%) to match the stiffness of ripe and rotten fruit, and established a two-choice oviposition assay within a four-quadrant circular arena to test the substrate hardness preference among four agarose concentrations with the hardness of 0.25% as control in ten *Drosophila* species (Figure 2B). The results revealed a clear behavioral difference. Most species displayed a strong and consistent preference for the softest substrate (0.25%) across all comparisons. Particularly, *D. mercatorum* and *D. biarmipes* that showed some oviposition success on real fresh fruit surface (Figure 1B-D), had strong aversion oviposition behaviors to hard agarose, suggesting that in nature, other cues like chemical cues might contribute to the override their innate mechanical avoidance of stiffness. However, *D. immigrans* and *D. suzukii* displayed quite distinct patterns from other species, and their preference for agarose were zero and showed no significant difference across 0.25% vs 0.5-1.5%. In some cases, the two fly species displayed slightly favoring the hard agarose of 1% and 1.5% (Figure 2B). These results indicate that *D. immigrans* and *D. suzukii* exhibit a relatively weaker preference for substrate hardness and have ability to lay their eggs on multiple fruit surfaces despite skin stiffness. It is possible to suggest the loss of oviposition behavior aversion to mechanically hard surfaces as the key behavioral traits that distinguished these two species from all others tested flies.

Next, we used no-choice assay to assess the oviposition rate of ten flies on different agarose concentrations (0.25% to 2.0%) (Figure 2C). The results revealed that most species that oviposited on soft substrates (0.25% and 0.5%) which mimicking decayed fruit decreased the egg-laying rates obviously at higher hardness concentrations (1.0% to 2.0%), but *D. suzukii* and *D. immigrans* maintained higher egg-laying rates even on the hardest agarose concentrations (1.0% and 1.5%), which mimic the stiffness of ripe fruit. Notably, *D. suzukii* exhibited the most robust oviposition performance, maintaining ovipositing rate even at 2.0% agarose (Figure 2C).

Additional, to assess the effects of different chemical cues on preference, we used the two-choice oviposition assay within a four-quadrant circular arena to test the preference of ripe or rotten fruit purees separately. In this assay, one-half of the arena was filled with a mixture of fruit purees (ripe or rotten) combined with 0.5% agarose, while the remaining half was occupied by 0.5% agarose alone (Figure S1). The results indicated that *D. suzukii* show an indifference oviposition preference on ripe strawberry puree and significant aversion to rotten strawberry puree, and *D. mercatorum* exhibited a slight preference for ripe strawberry puree and an intermediated preference on rotten strawberry puree, while other eight species, including *D. immigrans,* display a clear preference for egg-laying on both the ripe and rotten strawberry puree (Figure S1).

In summary, these results reveal the significant differences in oviposition behavior across *Drosophila* species. Both *D. suzukii* and *D. immigrans* possess oviposition capability to overcome the mechanical barrier of intact fruit surfaces or hard substrate. However, when exposed to chemical signals, *D. suzukii* exhibited highly specific responses, whereas *D. immigrans,* like most generalized species, was attracted to a broad spectrum of stimuli.

### Morphological specialized ovipositor facilitates the puncture oviposition to the fresh fruit with stiffness skin

Previous studies showed that *D. suzukii* and its sister species possess a distinctively long and serrated ovipositor with a capable of puncturing intact fruit skin (Atallah et al., 2014). Thus, we investigated the relationship between the morphology of ovipositor and the egg-laying position among the ten testing species. We categorized position of eggs deposited in strawberries and raspberries with hard or tough fruit surface into three types: “puncture” (egg inserted into intact skin), “embedded” (egg laid in a fruit crevice), and “surface” (egg deposited on the exterior) (Figure 3A).

**Figure 3.**
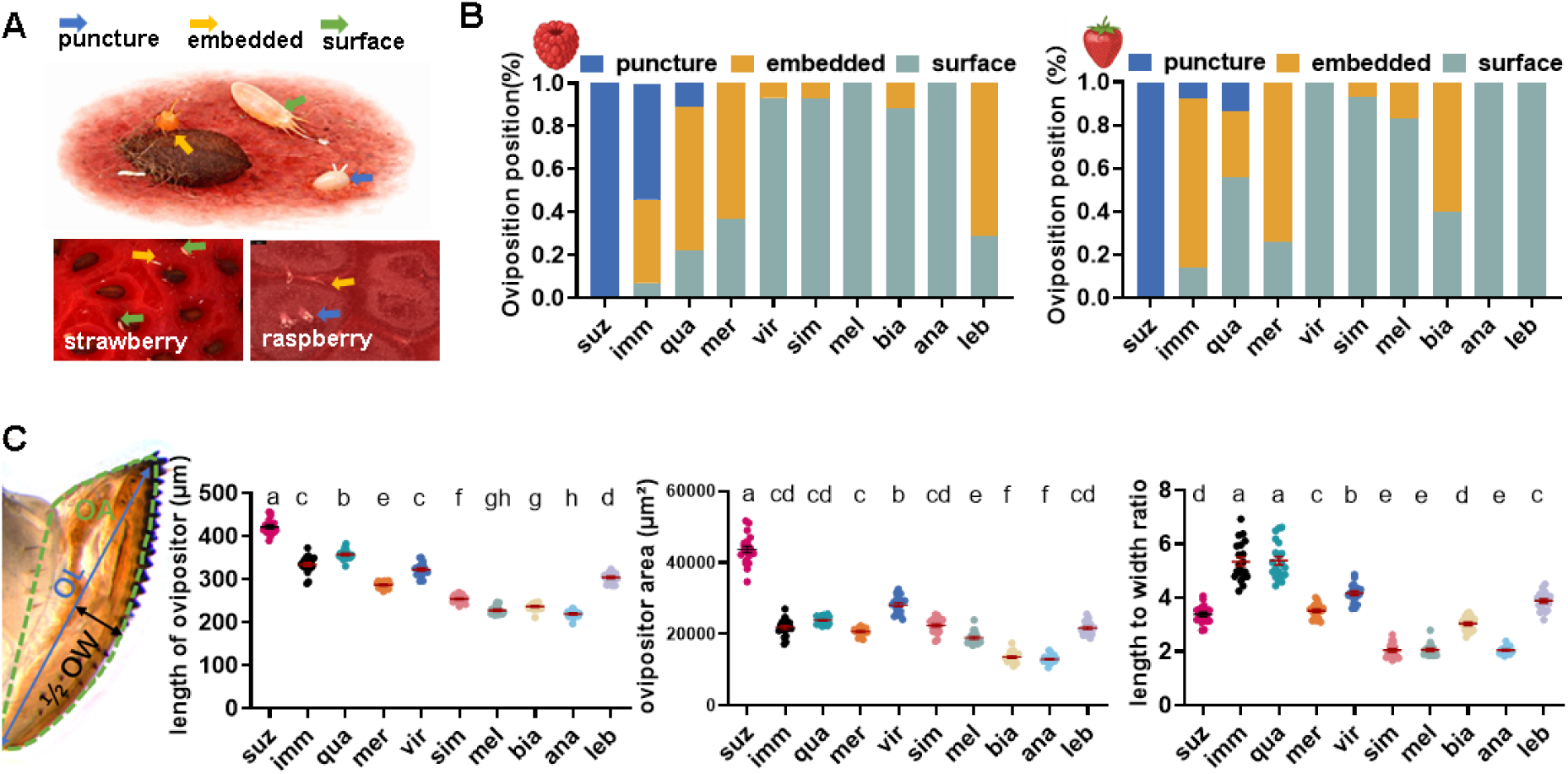
The ability of puncture oviposition in ripe fruit varied greatly among different species. (A) *Drosophila* show three types of oviposition position in fresh strawberry and raspberry. Yellow denotes Puncture Oviposition (PO), blue indicates Embedded Oviposition (EO), and green indicates Surface Oviposition (SO). (B) The oviposition position of ten species on ripe raspberry (i) and strawberry (ii). Blue denotes the proportion of PO; blue indicates the proportion of EO; green show the proportion of SO. Only *D. suzukii*, *D. immigrans*, and *D. quadrilineata* demonstrate the ability to puncture intact fruit skin. (C) Measurement standard of length, area and length to width ratio of ovipositor of *Drosophila* and morphological characteristics of ovipositors (left : length of ovipositor, middle: length to width ratio of ovipositor, right: ovipositor area). The data was analyzed by ordinary one-way ANOVA . The letters show that the two groups of data are significantly different (p < 0.05), n=20.

Five species were physically incapable of penetrating the fruit skin and were restricted to oviposit on the surface but *D suzuki* completely puncture the fruit skin and insert the eggs into the fruit (Figure 3B). The species of *D. immigrans* and its closely related species *D. quadrilineata* showed the unique puncture oviposition behavior and display three position types of egg deposition: puncture, embedded and surface (Figure 3B). We further used agarose substrate as control experiments to confirm the impact of different stiffness levels on oviposition behaviors of those two species. We found that *D. immigrans* and *D. quadrilineata* as well as *D. suzukii* are capable of puncture oviposition across all four agarose concentrations (0.25%, 0.5%, 1.0% and 1.5%). However, *D. mercatorum* showed a high overall oviposition rate on both hard agarose and ripe fruits, but only deposited eggs on the surface of agarose. This indicates that oviposition rate does not imply to puncture hard fruit skin (Figure 3B, STable).

Besides mechanical constraint, motivation can influence the oviposition on hard substrates. So, we supplemented the hardest agarose (2.0%) with a yeast–sucrose mixture to enhance nutritional motivation. This treatment markedly increased total egg laying on the surface of agarose across all ten fly species (Figure S2A) but failed to induce puncture oviposition in any species. Thus, nutritional motivation can restore the females’ oviposition but could not promote their ability to penetrate hard substrates. This confirms that the inability to puncture intact fruit skin is a mechanical constraint rather than a lack of nutritional motivation.

To determine the physical requirements for overcoming the mechanical defenses of intact fruit, we performed a comparative morphological analysis of the female ovipositors. We found that while most species of *Sophophora* subgenus, such as *D. melanogaster*, possess short, rounded ovipositors unsuitable for penetration, two distinct specialized forms have emerged among species capable of puncturing. In the specialist *D. suzukii*, the ovipositor is characterized by an exceptionally large surface area and a robust, ’saw-like’ structure (Figure 3C). In contrast, the species of *D. immigrans* in *Drosophila* subgenus have evolved a ’needle-like’ strategy, featuring significantly longer ovipositors with a high length-to-width ratio that results in a sharply tapered profile.

Together, these results show that the hard skin of ripe fruits inhibits both oviposition rate and puncture behavior in most *Drosophila* species. However, species like *D. suzukii*, and *D. immigrans* have unique morphological ovipositor. Generally, the elongated or serrated ovipositor can promote the *Drosophila* species to insert eggs through intact fruit skin during oviposition process. These associations suggest that ovipositor structure contributes to penetration capacity in *Drosophila*.

### The mechanosensory channel Inactive (*IAV*) contributes to stiffness-dependent inhibition of oviposition in saprophagous species *D. melanogaster*

Having established the importance of substrate stiffness cues in oviposition position of fruit flies, we asked whether mechanosensory channels of *Drosophila* flies are mediated their oviposition divergence on hard substrates. Among the genetic toolkit of model species *D. melanogaster*, there are eight mutants of mechanosensory ion channels including the DEG/ENaC channels *Pickpocket 1*(*ppk1*) (Gou et al., 2014) and *Pickpocket 26* (*ppk26*) (Guo et al., 2014), TRP channels *Nanchung* (*nan*) (Zhang et al., 2013), *Inactive* (*iav*) (Gong et al., 2004; Zhang et al., 2013), *NompC* (Yan et al., 2013), the evolutionarily conserved mechanosensitive channels *Piezo* (Kim et al., 2012) and *TMC* (Zhang et al., 2016), and the ionotropic receptor channel *Ir8a* (Bigge, 1999). For each mutant genotype, we checked the oviposition behavior and oviposition rate on four difference stiffness ranges (0.25%, 0.5%, 1.5% and 2.0% agarose), then compared them to the wild type *w^1118^*.

While several mutants (*nan^Gal4^*, *ir8a^1^*and *ppk26^Gal4^*) females exhibited generalized disinhibition on softer substrates, the loss of inactive (*iav*) mutant resulted in a distinct oviposition behavioral phenotype. Unlike wild type *w^1118^* controls, which exhibit a sharp decline in oviposition rate as stiffness increases, *iav^1^* mutants maintained high oviposition rates even on the stiffest substrates (1.5% and 2.0% agarose) (Figure 4A). Crucially, the loss of oviposition behavioral aversion was accompanied by a shift in behavioral pattern: *iav^1^* females successfully performed puncture oviposition on high-stiffness agarose, an oviposition behavior strictly absenting in wild-type *D*. *Melanogaster* (Figure S2B, STable). Moreover, *iav^1^* mutants only partially successful penetrated the intact skin, which quite differs from the complete penetration in the specialist species of *D. suzukii*. The *ir8a^1^* mutant also showed reduced oviposition behavioral aversion, but they failed to puncture intact fruit skin (strawberry and raspberry) (Figure 4B-D and STable).

**Figure 4.**
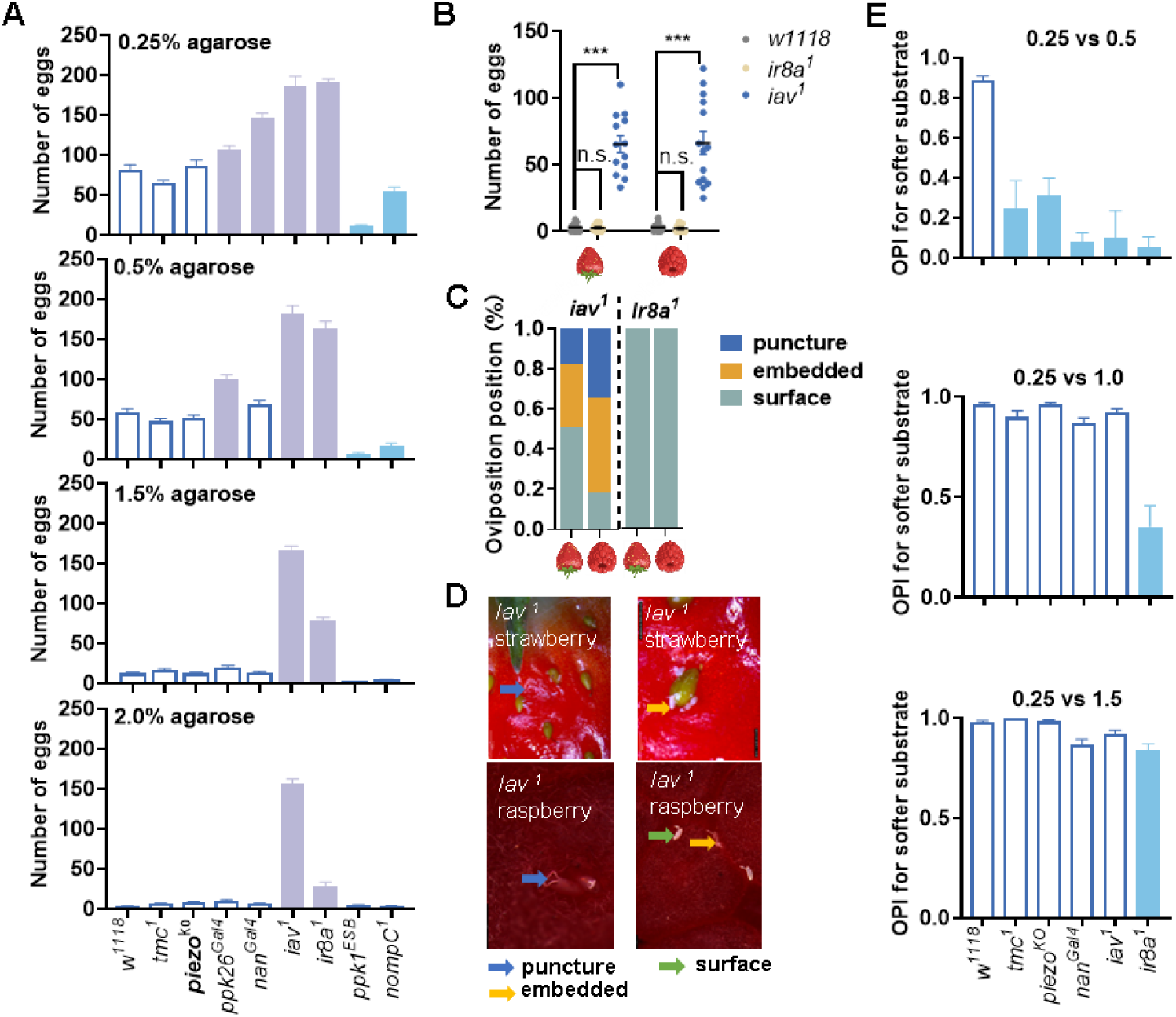
*Iav* is necessary for oviposition response to stiff substrates. (A) The number of eggs laid of *w^1118^* (*D. melanogaster*) and its eight mechanosensitive gene mutants at soft (0.25% and 0.5% agarose) and hard (1.5% and 2.0% agarose) substrates. *Iav^1^*mutants completely lose oviposition aversion on hard substrates, maintaining high oviposition rates. One-way ANOVA with Dunnett’s correction for multiple comparisons against the *w^1118^* group, Mutants with increased and decreased oviposition rate are highlighted in purple and blue. Error bars are SEM. n= 20-28. (B) The number of eggs laid of two mechanosensitive gene mutants (*iav^1^* and *ir8a^1^*) in whole ripe fruits (strawberry or raspberry) in a no-choice assay. Two-way ANOVA followed by Sidak’s multiple comparisons test. Error bars are SEM. n= 14-19. (C) The proportion of puncture oviposition by *iav^1^* and *Ir8a^1^* mutant females on hard substrates (ripe strawberry and ripe raspberry). Error bars are SEM. n=14-19. (D) *Iav^1^*mutant females puncture the intact skin of ripe fruit to egg-laying. Scale bars, 1 mm. (E) OPI for 0.25% of five mechanosensitive gene mutants at difference agarose ranges (0.5%, 1.0% and 1.5%). One-way ANOVA with Dunnett’s correction for multiple comparisons against the *w^1118^*group, Mutants with decreased oviposition rate are highlighted in blue. Error bars are SEM. n=12.

The species of *D. melanogaster* shows obviously preference to the softer substrates for oviposition (Figure 2B). Then, we compared the oviposition preference of eight mechanosensory mutants in stiffness substrate. Our results indicated that the mutants of *piezo^KO^*, *tmc^1^*, *nan^Gal4^*, *iav^1^* could discriminate the difference of 0.25% and 0.5% agarose, whereas *ir8a^1^* mutants impaired the discrimination for three stiffness levels (0.25 vs 0.5%, 1.0%, 1.5%), and all eight mutants showed the preference for softer substrates of 0.25% and 2.0% agarose (Figure 5E).

**Figure 5.**
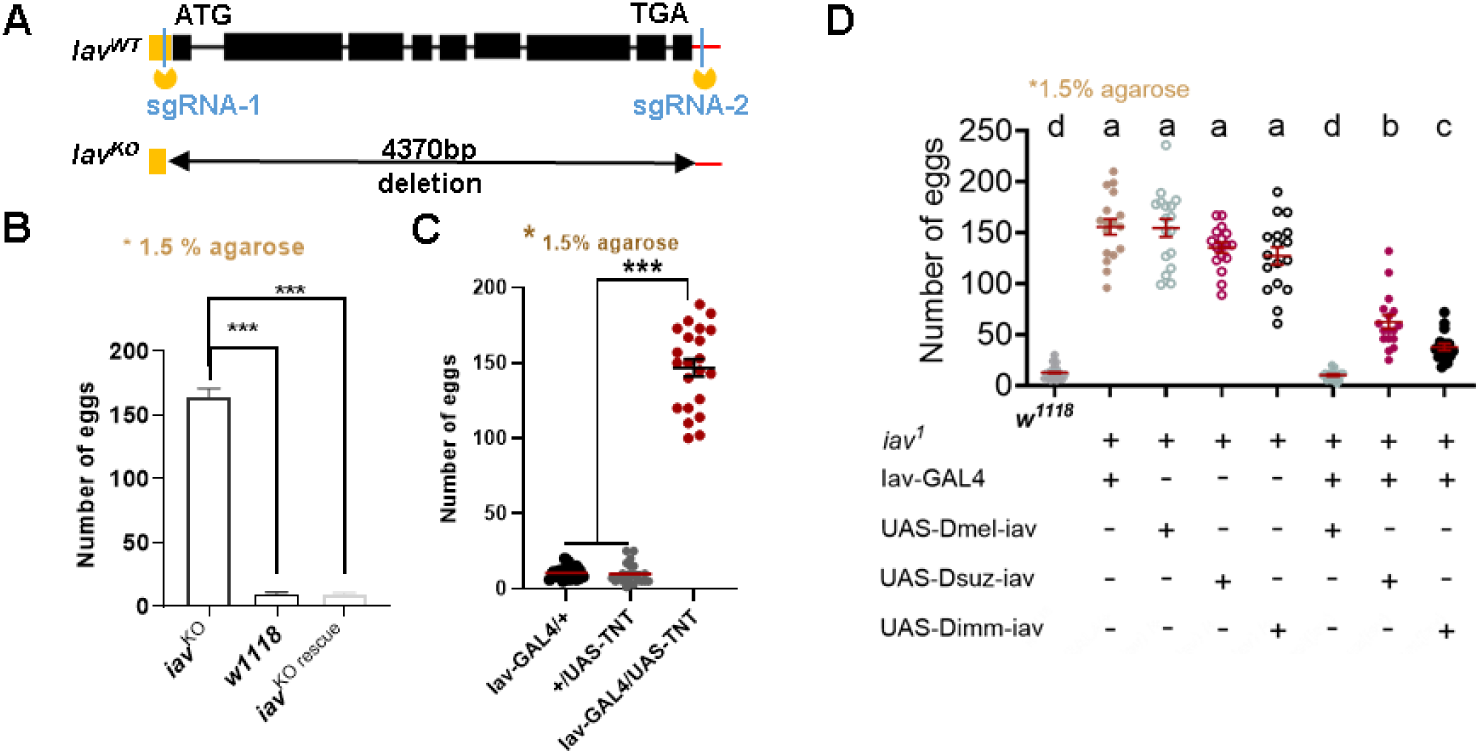
Neural activity of *Iav* expressing neurons is essential for oviposition response to stiff substrates. (A) Schematic representation of loss-of-function *Iav* obtained by CRISPR/Cas9-mediated mutagenesis in *D. melanogaster*. Knock out line was created using the sgRNAs indicated on the first line. (B) Behavioral responses of females of *Iav^KO^* mutants and transgenic rescue flies (Iav^KO^ ; Iav-GAL4; UAS-iav) to indicated substrate texture*. Iav^KO^* rescue type restores the innate inhibition of oviposition on stiff substrates. Error bars are SEM. n = 16. (C) The number of eggs laid on the 1.5% agarose when Iav-Gal4 neurons are blocked with TNT. Inactivation of iav neurons impairs the inhibition of oviposition on hardness substrate. one-way ANOVA with Dunnett’s correction for multiple comparisons against the Iav-TNT group. Error bars are SEM. n = 22. (D) The number of eggs laid on the 1.5% agarose. the Dsuz-iav and Dimm-iav rescues exhibited significantly higher oviposition rates than the Dmel-iav control, yet significantly lower rates than the *iav^1^* mutants. Genotypes were as follows: Iav^1^; Iav-GAL4/+ (GAL4 control), Iav^1^; UAS-Dmel(iav)/+ (UAS Dmel control), Iav^1^; UAS-Dmel(iav)/ Iav-GAL4 (GAL4>UAS Dmel rescue), Iav^1^; UAS-Dsuz(iav)/+ (UAS Dsuz control), iav^1^; UAS-Dsuz (iav)/ Iav-GAL4 (GAL4>UAS Dsuz rescue), Iav^1^; UAS-Dimm(iav)/+ (UAS Dimm control), iav^1^; UAS-Dimm (iav)/ Iav-GAL4 (GAL4>UAS Dimm rescue). Sgnificant differences are denoted by letters (two-way ANOVA followed by Tukey’s test for multiple comparison). Error bars are SEM. n = 17.

Together, our results identified the contribution of fruit flies’ mechanosensitive channel *iav* to puncture oviposition behavior in ripe fruits with stiffness skin. We also noted that the mechanosensory system regulates various aspects of oviposition behavior in female flies through multiple pathways, such as the *ir8a* has the function in oviposition preference of substrate stiffness.

### *IAV* orthologs differ in their restoration of the innate inhibition of stiffness-dependent oviposition

To further validate the role of *iav* in sensing substrate stiffness, we generated a null *D. melanogaster* mutant *iav* knockout (KO) line *iav^KO^* using CRISPR-Cas9 (Figure 5A). Consistent with *iav^1^* mutant, the *iav^KO^* lines exhibited the complete loss of oviposition behavioral aversion to hard substrates, but maintained the high oviposition rates on 1.5% agarose compared to wild type (Figure 5B). This sensory deficiency can be fully rescued by expressing a *D. melanogaster UAS-Dmel-iav* transgene under the control of the *iav-Gal4* driver, which restored the innate inhibition of oviposition on stiff substrates (Figure 5B and 5D). Furthermore, blocking synaptic transmission in *Iav-Gal4* neurons using tetanus toxin light chain (TNT) significantly impaired the inhibition ability of oviposition on 1.5% agarose (Figure 5C). However, the neurons silencing oviposition behavioral phenotype on hard agarose had less inhibition ability than that of the *iav^1^* mutant in ripe strawberry because the late showed almost no puncture oviposition in the ripe strawberry. This result suggests that the *iav* channel and its associated sensory neurons (Iav-expressing) are critical for texture detection, although the Iav-Gal4 driver may not capture the complete expression pattern of the endogenous gene in the mechanosensory system.

Given that *D. suzukii* and *D. immigrans* showed a weaker oviposition inhibition on stiff substrates than *D. melanogaster* (Figure 2C), we hypothesized that the *IAV* channel itself might have functionally diverged that contributes to this behavior change. Sequence comparison revealed notable amino acid variations between the orthologs of these species of *D. suzukii* and *D. immigrans*, and *D. melanogaster* (Figure S3). To test whether these variations contribute to species-specific behaviors, we cloned the *iav orthologs* from *D. suzukii* and *D. immigrns* and introduced them into the *D. melanogaster iav^1^* mutant background. We constructed two UAS-line (UAS-Dsuz-iav and UAS-Dimm-iav) and tested behavioral responses to stiff substrate in animals in which the UAS-line-iav transgene was the only source of *iav*.

Both heterologous rescued lines (*Iav^1^*; Iav-Gal4/Dsuz-iav and Iav^1^; Iav-Gal4/Dimm-iav) retained the ability of puncture oviposition on 1.5% agarose, and significantly exceeding the oviposition rates of the Dmel-iav rescue while remaining below the total disinhibition in *iav^1^* mutants (Figure 5D). Those results demonstrated that the donor species’ *iav* ortholog is partially recapitulates their specialized behavioral phenotypes (stiffness tolerance) within the physiological context of *D. melanogaster*. Thus, our results for the first time provide direct evidence that the *IAV* gene has functionally diverged across these lineages, and that modifications to this mechanosensory channel are a key molecular mechanism allowing species to overcome the physical defenses of their host plant.

### Niche overlap and natural resource differentiation among co-occurring *Drosophila* species

An oviposition strategy is adaptive only if it confers a survival advantage to the offspring (Kershenbaum et al., 2012). To connect oviposition behavior with fitness, we measured survival rate of the offspring after oviposition assay. We found that the ability to puncture or embed eggs in fresh fruit significantly impacts larval success. In fresh strawberries, only the offspring of *D. suzukii* and *D. immigrans*, the species capable of penetrating the intact skin consistently reached adulthood (Figure 6A). Notably, the puncture behavior has obviously initiated or accelerated the localized fruit decay. By the third day, strawberries treated with *D. suzukii* or *D. immigrans* exhibited visible breakdown, providing larvae with essential access to softened tissue and nutrients. By contrast, fruit exposed to *D. melanogaster* showed minimal decay on the third day after oviposition assay, leaving larvae unable to penetrate the substrate or acquire sufficient nutrition (Figure 6B). Conversely, on soft skinned raspberries, all ten species larvae developed successfully into adults, as the fruit naturally decayed within 2–3 days regardless of the oviposition method (Figure 6A).

**Figure 6.**
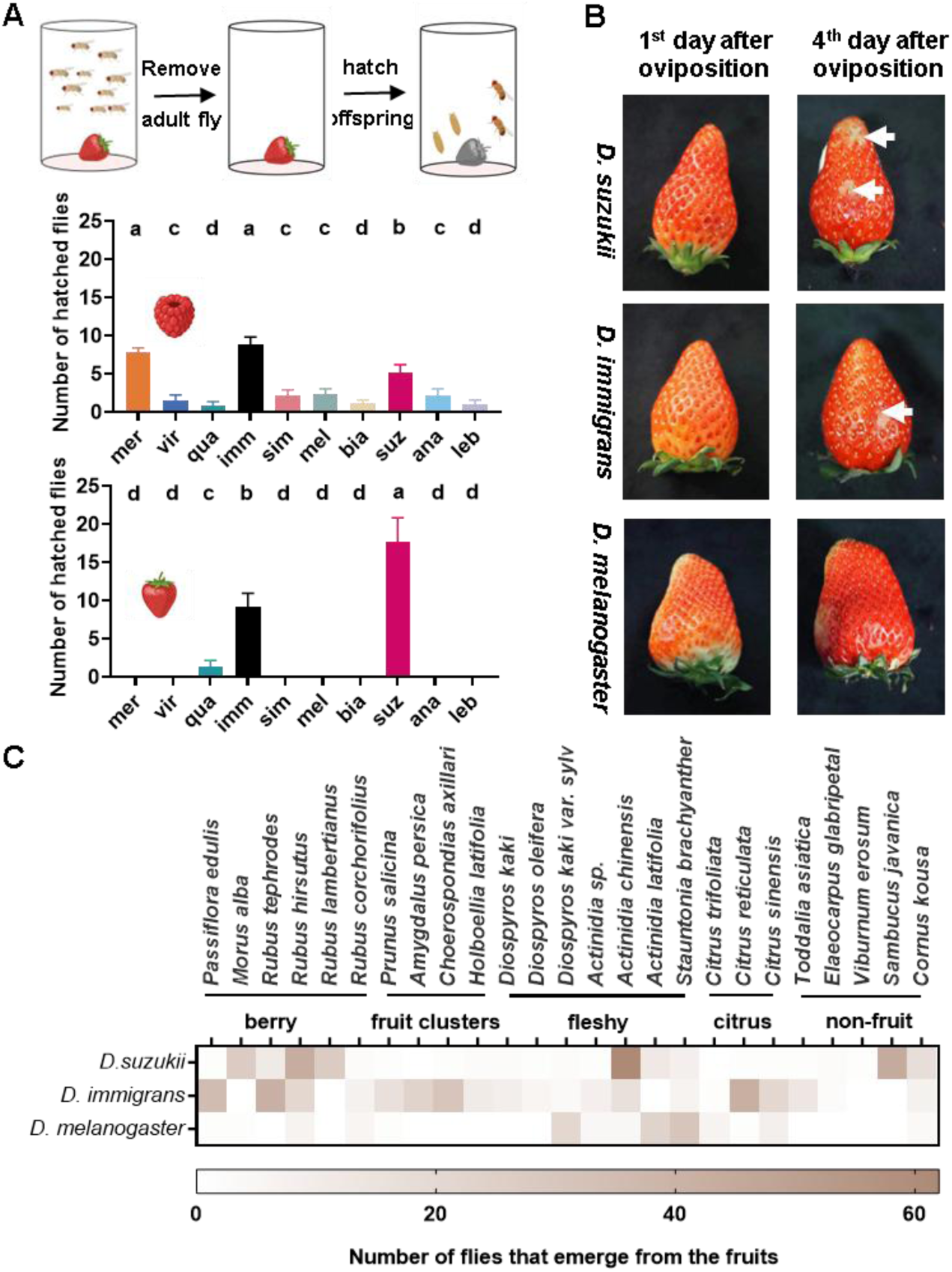
*D. immigrans* and *D. suzukii* utilize similar ecological niches. (A) Number of hatched flies on whole ripe raspberry (left) and strawberry (right) in a no-choice oviposition assay. Only species capable of penetrating the intact skin (*D. suzukii* and *D. immigrans*) successfully complete larval development in fresh strawberries. Significant differences between species are denoted by letters (p<0.05). Two-way ANOVA followed by Tukey’s test for multiple comparison). Error bars are SEM. n=12. (B) Differences in the decaying condition of fruits between 1st and 4th day after oviposition by three species females. The fly species and days after oviposition are labeled in white text. Both *D. suzukii* and *D. immigrans* experiments have obvserved the apparent strawberry decay on the 4th, while there was almost no change in *D. melanogaster* treatment. (C) Hatching number of *D. immigrans*、*D, melanogaster and D. suzukii* from natural substrates in Shunhuang Mountain.

To determine if above laboratory findings can indicate the circumstance under natural field conditions, we conducted a comprehensive field survey to understand the egg-laying site distribution of *Drosophila* species in their natural habitat. We collected local fruit fly samples from the Shunhuang Mountain region (Hunan province, the central China, a critical biogeographical transition zone representing the ancestral origins of many core *Drosophila* groups (Robert Liu et al., 2015)). We systematically collected potentially substrate about 24 plant species, including various fresh fruits, flowers, and rotting plant materials. These field collected samples were subsequently incubated in the laboratory, and the emerging adults were reared to identify the specific species capable of completing their development within each host. By focusing on rearing larvae to adulthood from collected fruit, we ensured that our survey accurately reflected the reproductive success and ecological niche occupancy of each species under natural conditions (Figure 6C). Results showed that the community was consistently dominated by three species (*D. melanogaster*, *D. immigrans*, and *D. suzukii*) across nearly all fruit samples. Despite the species of *D. suzukii* is phylogenetically closer to *D. melanogaster,* its local egg-laying distributions of eggs were similar to that of *D. immigrans*. Both *D. suzukii* and *D. immigrans* species were frequently co-occurred on the same sets of firm-surface fruits, such as the fruits of *Rubus* and *Actinidia* species. Furthermore, they have a broader distribution range than *D. melanogaster*. These findings demonstrate that the ability to overcome mechanical defenses facilitates their ecological adaptability, mediating a significant expansion of their ecological niche in natural habits.

## DISCUSSION

### Overcoming mechanical barriers contributes to niche breadth

Niche expansion often proceeds through the colonization of resources protected by physical or chemical barriers (Sexton et al., 2017). Most *Drosophila* species are restricted to a saprophagous niche, relying on the successional decay of fruit to provide a soft substrate for larval development (Markow and O’grady, 2008). While fruit ripening is a complex process involving concurrent changes in both chemistry and physical texture, our findings clarify the relative importance of these constraints. Ecological theory distinguishes substitutable from non-substitutable resources, showing that niche structure differs fundamentally depending on whether one resource axis can compensate for another (Ashby et al., 2017). Chemical properties of fruit are relatively substitutable: insects can evolve shifts in sensory tuning to track different volatile profiles across varying decay states. By contrast, our agarose decoupling assays demonstrate that the mechanical hardness of intact fruit skin functions as a strict, non-substitutable barrier (Figures 1 and 2). For many generalist species, avoidance of firm substrates is not a result of reduced chemical attraction, but a mechanical exclusion. In this framework, *D. suzukii* and *D. immigrans* have each overcome this barrier to access the ripe-fruit niche, but the ecological consequences of doing so are different: *D. suzukii* largely abandons the ancestral rotten-fruit niche, representing niche specialization, whereas *D. immigrans* retains broad use of decaying substrates alongside its newly gained access to fresh fruit, representing true niche breadth expansion.

Crossing a mechanical barrier requires both behavioral intent and physical tools (Crava et al., 2020; Goldman-Huertas et al., 2015; Karageorgi et al., 2017). Previous work established that *D. suzukii* depends on its enlarged, saw-like ovipositor to puncture intact fruit skin, and that this morphological specialization is central to its ecological shift (Karageorgi et al., 2017). Our comparative morphological analysis reveals a second independently evolved solution: *D. immigrans* possesses a needle-like ovipositor with a high length-to-width ratio and a sharply tapered tip that enables penetration through the skin (Figure 3C).These two morphologies represent convergent mechanical strategies, each adapted to breach the same physical barrier through distinct structural principles. Crucially, the ecological significance of puncture oviposition extends beyond the act of egg deposition itself. Penetration of fresh fruit skin by *D. immigrans* initiates localized decay, providing developing larvae with access to softened tissue and nutrients that remain inaccessible to species restricted to surface oviposition (Figure 6A). The close association between puncture behavior and larval survival on firm hosts confirms that morphological innovation translates directly into a fitness advantage for the two species.

### Neural Mechanosensory Solutions to the Stiffness Barrier

Possessing an ovipositor capable of penetrating a firm surface is unlikely to be sufficient for exploiting intact fruit if the nervous system strongly suppresses oviposition on stiff substrates. In generalist species like *D. melanogaster*, encountering high substrate stiffness triggers a mechanosensory constraint that strongly inhibits oviposition (Zhang et al., 2020). To understand the neural basis of overcoming this barrier, we screened a series of mechanosensory mutants and identified the Inactive (IAV) ion channel as an important regulator of stiffness dependent oviposition inhibition in *D. melanogaster*. (Figure 4). Unlike wild-type *D. melanogaster*, which exhibit a sharp decline in egg-laying on firm substrates, *iav* loss-of-function mutants maintained high oviposition rates on the stiffest media (Figure 4A and 5B). Crucially, removing this sensory constraint unlocked a latent behavioral capacity, *iav* mutants successfully performed puncture oviposition on high-stiffness agarose, a behavior strictly absent in wild type *D. melanogaster* (Figure 5D). Nevertheless, the limited penetration achieved by iav mutants shows that removing this inhibition is not sufficient to result the complete fruit-piercing ability of *D. suzukii* or *D. immigrans*. Effective exploitation of intact fruit therefore likely requires both reduced behavioral inhibition and appropriate special morphological ovipositor. It is important to note that the mechanosensory system regulates oviposition through multiple, complex pathways. For instance, while mutants for the ionotropic receptor *Ir8a* also showed reduced behavioral aversion and impaired stiffness discrimination (Figure 4A and 4E), they entirely failed to puncture intact fruit skin. These results suggest that substrate discrimination, behavioral inhibition, and physical penetration are associated but separated components of the whole oviposition process. Within this framework, *IAV* contributes to the mechanosensory regulation of oviposition puncture behavior, but it should not be considered the sole determinant or direct mediator of this mechanical penetration. Adapting to a new niche commonly includes conserved pathways being recalibrated rather than the complete loss of an essential sensory channel (Auer et al., 2020; Goldman-Huertas et al., 2015; Karageorgi et al., 2017). Given that both *D. suzukii* and *D. immigrans* show reduced oviposition inhibition on stiff substrates relative to *D. melanogaster*, we asked whether their *IAV* orthologs differed functionally under a common genetic background. Sequence comparison revealed notable amino acid variations between the *IAV* orthologs of *D. suzukii* and *D. immigrans* comparing to that of *D. melanogaster* (Figure S3). We therefore expressed the *D. suzukii* and *D. immigrans* orthologs in the *D. melanogaster iav* mutant background and compared their ability to restore stiffness dependent oviposition inhibition. Both heterologous rescue lines retained the ability to perform puncture oviposition on 1.5% agarose and showed significantly higher oviposition rates than the *D. melanogaster IAV* rescue, while remaining below the complete disinhibition seen in *iav* null mutants (Figure 5D). The cross-species rescue experiments further show that the *IAV* orthologs differ in their ability to restore stiffness-dependent oviposition inhibition under a common *D. melanogaster* background. This result is consistent with functional divergence among the orthologs of those three *Drosophila* species. However, because the currently experiments were conducted in a heterologous genetic and cellular context, they can not directly demonstrate that IAV divergence results the natural behavioral or ecological niches diversity among *D. melanogaster, D. suzukii*, and *D. immigrans*. Additional experiments in the native species, together with physiological measurements of channel function and expression, would be required to establish such an evolutionary niche contribution.

### Evolutionary Strategies and Ecological Partitioning in *Drosophila*

The diversification of ecological niches in the genus *Drosophila* reveals multiple evolutionary routes to novel resource use (Chialvo et al., 2019). A central distinction is whether a lineage gains access to a new resource while retaining its ancestral one, thereby expanding niche breadth, or instead gains the new resource by abandoning the ancestral niche, thereby becoming more specialized (Sexton et al., 2017). In this framework, our comparison across *Drosophila* species highlights three distinct evolutionary routes to niche exploitation. First, *D. sechellia* represents a classic case of niche specialization driven largely by chemosensory evolution. It lays eggs almost exclusively on noni fruit, and this behavioral shift depends strongly on evolved olfactory and taste system, including *Ir75b*-mediated responses to host-specific odorants (Dworkin and Jones, 2009; Kadow and Gompel, 2020) (Álvarez-Ocaña et al., 2023; Auer et al., 2020). *D. suzukii* represents a different specialist route. It evolved the ability to use ripe fruit through a serrated ovipositor, altered mechanosensory responses, and chemosensory changes associated with ripe-fruit oviposition. However, this gain was coupled to a narrowing of the ancestral saprophagous niche, resulting in a reduced use of rotten substrates and aversion to late fermentation cues (Figure S1)(Keesey et al., 2015). By contrast, *D. immigrans* illustrates niche expansion in stricter sense. It possesses an elongated, needle-like ovipositor that provides mechanical access to firm substrates, while retaining broad use of rotting substrates and other ancestral substrates, thereby widening rather than replacing its original resource range (Figure 3 and S1).

This contrast becomes clearer when the ripe-fruit niche is viewed as a combination of separable chemical and mechanical axes. Along the fruit-ripening continuum, volatile composition, sugar and acid balance, and surface stiffness all change, but these dimensions do not impose equivalent ecological constraints. Chemical sensory system exhibit high evolutionary plasticity, characterized by functional shifts in receptors, differential expression patterns, and olfactory sensory neuron abundance (Takagi et al., 2024; Zhao and McBride, 2020). These modifications are frequently coupled with niche specialization, enabling the species to successfully explore novel ecological environments (Zhao & McBridge, 2020; Hansson & Stensmyr, 2011; Ramdya & Benton, 2010). However, mechanical hardness is different. In ecological terms, the intact skin of ripe fruit can function as a non-substitutable barrier. A female may detect and even be attracted to fresh-fruit chemical signals, yet still fail to use that resource if they cannot pierce the oviposition site (Keesey et al., 2015). Our comparison with *D. biarmipes* is consistent with this view. Although this species can show attraction to ripe fruit, it remains strongly limited by stiffness and fails to convert attraction into effective use of firm fruits (Figure 3B and S1). Therefore, the key step in accessing ripe fruit is not simply a change in preference, but rather the acquisition of a mechanical solution that overcomes a barrier that other sensory cues cannot bypass.

Previous work in Drosophilids has shown that narrow ecological specialization often arises through coordinated changes across multiple organs and sensory systems. The genus provides an especially powerful comparative framework because the genetic tools developed in *D. melanogaster* can be extended to related species occupying distinct ecological niches (Auer et al., 2020). In *D. sechellia*, host specialization involves evolved olfactory tuning, altered sensory circuit organization, and gustatory modifications associated with toxic-fruit use (Takagi et al. 2024; Prieto-Godino et al., 2017; Auer et al., 2020). In *D. suzukii*, the shift toward ripe fruit likewise reflects coordinated sensory evolution, including altered responses to fermentation-associated cues, changes in sweet valuation, and modifications in stiffness-related oviposition behavior (Figure S1) (Cavey et al. 2023; Wang et al. 2022). Together, these cases suggest that niche specialization often depends on multisensory recalibration rather than on a single trait. Our results indicate, however, that niche expansion can follow a different logic. *D. immigrans* does not appear to achieve access to fresh fruit by narrowing its ancestral chemical niche. Instead, its oviposition responses remain broadly overlapping with those of more generalist species (Figure S1). Its elongated, “needle-like” ovipositor and reduced inhibition on stiff substrates provide the mechanical capacity to exploit a previously unsuitable resource (Figure 3C and S3). It is this combination that makes the *D. immigrans* strategy exceptional. Rather than replacing ancestral decaying-fruit preference with a fresh-fruit preference as a specialist, *D. immigrans* retains a generalist chemical background and adds a mechanical innovation to it, thereby extending its resources use across the full ripening gradient, from fresh to rotten fruit.

The ecological consequences of this difference are evident in the field. Our comprehensive field survey in the Shunhuang Mountain region validates that mechanical capability acts as an ecological filter structuring wild *Drosophila* communities (Figure 6C). Despite *D. suzukii* and *D. melanogaster* being more closely related phylogenetically, the local distribution of *D. suzukii* mirrored that of the more distantly related *D. immigrans*. Both species consistently co-dominated firm-surfaced hosts, while *D. melanogaster* was largely excluded from these resources. This striking niche overlap demonstrates that the convergent ability to overcome mechanical defenses directly translates to ecological partitioning in the wild, allowing these species to access a protected resource pool and reduce direct competition with generalist populations.

## MATERIALS and METHODS

### Drosophila stock

All flies were reared on standard cornmeal medium at 24°C, 60% relative humidity, on a 12-hour light-dark cycle (lights on at 8:00am). Egg-laying experiments were conducted under the same conditions. Standard cornmeal medium consisted of, per 1L, 6 g agar, 7.5 g sucrose, 50 g maltodextrin, 24.5 g yeast, 73 g corn meal, 17.5ml methyl 4-hydroxybenzoate, 4ml propionic acid.

For behavioral assays we used a set a wild type strains from Ehime Stock Center (ESC) included: *D. melanogaster* (*w^1118^*), *D. biarmipes* (CJB214), *D. suzukii* (HJ), *D. meractorum* (mer1527.01), *D. virilis* (viri-HUE), *D. quadrilineata* (quad-MTK), *D. immigrans* (MT91-1), *D. simulans* (Rakujuen), *D. ananassae* (AABBg1), *Scaptodrosophila lebanonensis* (leb0011.00).

Transgenic lines from Bloomington Drosophila Stock Center (BDSC) included: *nan^Gal4^* (68205), *piezo^KO^* (58770), *tmc1* (66556), *iav-Gal4 III* (52273), *UAS-TNT II* (28837), *ir8a^1^*(5909), *nompC^1^* (42260) and *iav^1^* were kindly provided by Dr. Yuh-Nung Jan UCSF, *ppk26^Gal4^* was kindly provided by Dr. Zuo-Ren Wang CAS, *ppk1^ESB^* was kindly provided by Dr. Benjamin A. Eatono THSC. Mutants used in this study have been backcrossed into *w^1118^* background.

### Egg-laying assays

For all oviposition assays, flies were collected on the first day of emergence and aged for 7-8 days in food vials with standard cornmeal medium. 4-5 days prior to the experiment, males and females were placed in a new food vial supplemented with high nutrition medium (1L ddH2O +65g yeast + 65g sucrose) for mating and producing eggs. For each trial, flies were placed into each chamber through ice anesthesia, and left 12 or 24 hours. Eggs were counted from each substrate. The specifics of each experiment are given below.

### Two-choice oviposition assay on fruits

For each trial 20 females were placed in a columnar experimental cage (height 10cm, diameter 10cm) which contained both ripe strawberry and rotten strawberry. Trials last 12 hours and are carried out in the dark, n= 12 replicates for each fly species. Eggs were counted from each fruit, and an oviposition preference index was calculated as follows: (number of eggs on ripe fruit – number of eggs on rotten fruit) / (number of eggs on ripe fruit + number of eggs on rotten fruit). The ripe strawberries with intact skin were purchased from a supermarket at the day of the experiment (Carefully examined under a stereoscope and only skin intact fruits were selected), and rotten strawberries (same variety) were allowed to rot in a 24°C and 60% relative humidity incubator for five days prior to the experiment (Check if the fruit has visible bruises, soft spots, or discoloration on the surface).

### No-choice oviposition assay on ripe fruits

For each trial, 10 females were placed in a columnar chamber (height 10cm, diameter 8cm) which contained one ripe fruit with intact skin (strawberry, raspberry or blueberry). Trials lasted 12 hours and were carried out in the dark, n= 12-19 replicates for each species. Eggs were counted according to the oviposition position. Ripe strawberries and blueberries were purchased from a local supermarket, and raspberry were picked from the orchard of the botanical experiment gardens at the day of the experiment.

### No-choice oviposition assay for substrate stiffness

Plastic Petri dishes without quadrant dividers (diameter: 90mm) were used for egg-laying chamber. Experimental substrates contained increasing concentration (0.25%, 0.5%, 1.0%, 1.5%, 2.0%) of agarose. Trials lasted 24 hours and were carried out in the dark, using 10 females per replicate (n=18-22).

### No-choice oviposition assay for stiff substrate with nutrition

Plastic Petri dishes without quadrant dividers (diameter: 90mm) were used for egg-laying chamber. Experimental substrates contained with a mixture of 6.5g yeast and 6.5g sucrose added to 2.0% agarose. Trials lasted 24 hours and were carried out in the dark, using 10 females per replicate (n=18-20).

### Two-choice oviposition assay on substrate

Plastic Petri dishes (Nast, China) with four-quadrant dividers (diameter: 90mm) were used for egg-laying chamber. Quadrants were alternatingly loaded with 5 mL agarose in specific concentrations. Unless otherwise indicated, agarose solutions were not supplemented with any other chemicals. The filled petri dishes were allowed to solidify for 1 hour at room temperature. 20 females were separately placed on an ice pad and introduced immediately into the egg-laying chamber. Flies were allowed to lay eggs for 24 hours, after which the total number of eggs laid on each agarose pad was counted, and the oviposition preference index was calculated as follows: (number of eggs on experimental substrate – number of eggs on control substrate)/ (number of eggs on experimental substrate + number of eggs on control substrate). Only assays where flies had laid more than 20 eggs were included in the final analyses, and 12-15 replicate assays were performed for each fly species at each concentration point. For stiffness preference assay, experimental substrates contained increasing concentration (0.5%, 1.0%, 1.5%) of agarose; control substrates were 0.25% agarose. For strawberry puree assay, experimental substrates were filled with 35% w/v strawberry puree (ripe or rotten) in 0.5% agarose. Ripe strawberries puree was made of mechanical disruption of whole ripe fruits at the day of the experiment, and the rotten strawberries puree was made from the ripe strawberries puree that allowed to rot in a 24°C and 60% relative humidity incubator for 4-5 days prior to the experiment. The strawberry puree was mixed together with molten agarose which cooled about 45-50°C. For preference assay between ripe and rotten strawberry puree, experimental substrates were filled with 35% w/v ripe strawberry puree and 35%w/v rotten strawberry puree in 0.5% agarose. For ripe or rotten strawberry puree preference assay, experimental substrates were 0.5% agarose and contained 35%w/v ripe or rotten strawberry puree, control substrates were 0.5% agarose.

### Oviposition substrate stiffness hardness measurements

We used the Semmes–Weinstein Monofilament set (Von Frey, Germany) to compare the stiffness of substrates of different agarose concentration and fruits. Agarose plates were prepared from a 20ml solution in a 90mm diameter dish (Nast, China). Measurements were following the procedure described in (Sánchez et al., 2017).

### Dietary substrate survey of fruit flies

Fieldwork involved systematic sampling of a broad range of potential fruit fly substrates across elevations, including diverse fruits, flowers, fungi, mosses, and humus. All collected substrates were placed in custom-designed emergence chambers for incubation. Emerged adult flies were collected daily and preserved in 75% ethanol. Species identification was performed using morphological examination supplemented with DNA barcoding.

### Generation of transgenic reagent

Genomic fragments including *iav* were PCR cloned from *D. melanogaster*, *D. immigrans* and *D. suzukii* with the primers P1 and P2 (species), and cloned into the pJFRC28–10 ×UAS-IVS-GFP-p10 transformation vector to generate the combination vector UAS-iav. These plasmids were microinjected into the germ-line of *w^1118^ iav ^1^* strain and integrated into the attP40 site on the second chromosome through phiC31 mediated gene integration by Fungene Biotechnology (http://www.fungene.tech). Then, three independent UAS-iav transgenic line was obtained and balanced.

Primer Sequences following, P1: GCAACTACTGAAATCTGCCA; P2 (mel): GCAACTACTGAAATCTGCCA; P2 (imm): CAGGCCTTGTGATCCTCCCACTT; P2 (suz): CAGATACTCCTCGCCCTCCATCA).

### *Iav sgRNA* design and synthesis

We targeted the following CG4536 sequences: CCAGGACAACGAGCTGGCTTCCT, CTAAGTGCTAGGTTTAAAAGTGG. Oligos were used to synthesize in vitro the two sgRNA with T7 RNA polymerase: sgRNA-1 was coupled with the oligo sgRNA-2. The two sgRNAs and their corresponding single-stranded oligos (ssODN) were all coinjected, together with Cas9 protein, at the following concentrations: 50 ng/µl of each sgRNA, 300 mg/µl of protein and 125 ng/µl of each single-stranded DNA oligo. Adults and their progeny were screened by PCR on genomic DNA extracted from single legs or entire individuals (single flies or pairs of flies) to detect deletions of single stranded DNA oligos. Hits were confirmed by Sanger sequencing of the PCR product. Primers used for screening were: Forward-CCGATACTCTAGCACTAAGTTATA; Reward-GATGGCGCAATCAATGCCCGCT

### Imaging

Images of fruit infested with eggs were taken under a microscope (Leica DM6B, Germany). Images of fruits were taken under a camera (Olympus E-M1Marklll, Japan).

### Quantification and statistical analysis

We used the GraphPad Prism 8 software package to graph and statistically analyze data. The normal distribution of data was tested by Kolmogorov-Smirnov test before performing statistical analysis. When data were normally distributed, we used parametric tests; when data were not normally distributed, we used non-parametric tests. All data are presented as mean ± s.e.m. We also consistently labeled the sample numbers directly on graphs.

## ACKNOWLEDGMENTS

We thank Dr. Wei Zhang at Tsinghua University for providing fly lines and Dr. Lisha Shao at Delaware University for comments on manuscript revisions. This work was funded by the National Natural Science Foundation of China (31670231, 31470278 and 31270430) to Y. L. and (31700198) to B. L.

## AUTHHOR CONTRIBUTIONS

Y.B.L. and B.Y.L. conceived the project and designed the study. S.H, Y.J.Y., X.N.Y. and W.F.Z. performed the collected the samples and performed behavior experiments. S.H., and T.W. wrote and revised the manuscript. T.W., B.Y.L. and Y.B.L. assisted with the data analysis, results interpretation, and manuscript revision. All authors read and approved the final manuscript.

