## Supplementary Material for "Ovipositor morphology and mechanosensory innovation traits favor niche width expansion in *Drosophila*"

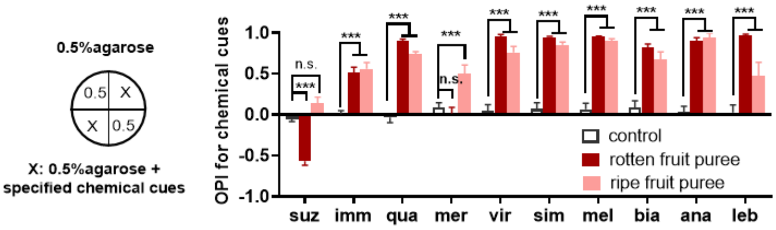


**Supplementary Figure 1. Oviposition preference for substrates associated with fruit maturation differ among 10 species.** Oviposition preference index of ten focal species for chemical cues between ripe fruit and rotten fruit. The fruit puree was used to present the chemical cues. 20 *Drosophila* females had the choice between two substrates of equal stiffness (0.5% agarose) but different chemical cues. Quadrants were alternatingly loaded with ripe and rotten strawberry puree. P-values were calculated using a Wilcoxon signed rank test against a theoretical value of 0 (no preference). Error bars are SEM. n =12.

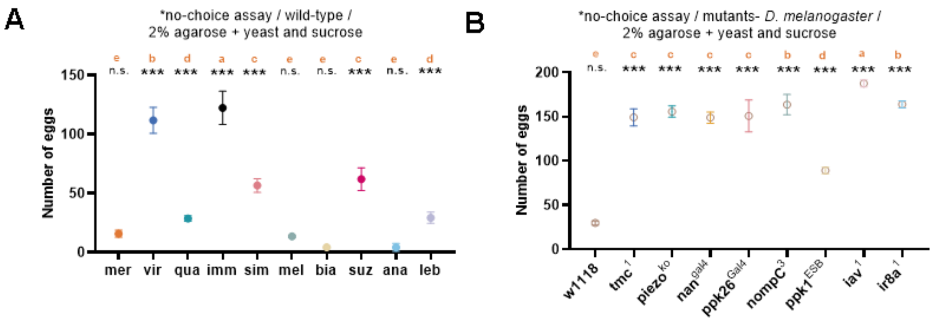


**Supplementary Figure 2. Substrate stiffness cues are related to oviposition position.**

(A) The oviposition rate of ten fly species on hard substrate (2.0% agarose) containing of yeast and sucrose. P-value was compared to 2.0%agarose using the Wilcoxon signed-rank test; significant differences between species are denoted by letters (two-way ANOVA followed by Tukey's test for multiple comparison). Error bars are SEM. n=18-20.

(B) The oviposition rate of six mechanosensory mutants and $w^{1118}$ on hard substrate (2.0% agarose) containing of yeast and sucrose. P-value was compared to 2.0% agarose using the Wilcoxon signed-rank test; significant differences between species are denoted by letters (two-way ANOVA followed by Tukey's test for multiple comparison). Error bars are SEM. n=12-16.

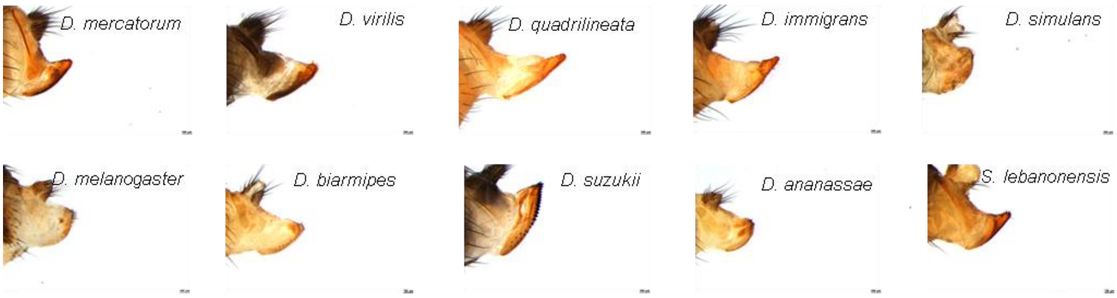


**Supplementary Figure 3. Ovipositor morphology in 10 *Drosophila* species from three subgenera.** Representative images highlight morphological variations among species belonging to three different subgenera. Scale bars, 100 um.

**Supplementary Table. The influence of substrate stiffness and nutrition in oviposition position.**

| Wild-type | Substrate | Oviposition position |
|---|---|---|
| mer | 0.5% agarose / S1 | P |
| | 2.0% agarose / S2 | S |
| | 2.0% agarose + yeast and sucrose / S3 | S |
| vir | S1 | P |
| | S2 | S |
| | S3 | S |
| qua | S1 | P |
| | S2 | P |
| | S3 | P |
| imm | S1 | P |
| | S2 | P |
| | S3 | P |
| sim | S1 | P |
| | S2 | S |
| | S3 | P |
| mel | S1 | P |
| | S2 | S |
| | S3 | S |
| bia | S1 | P |
| | S2 | S |
| | S3 | S |
| suz | S1 | P |
| | S2 | P |
| | S3 | P |
| ana | S1 | P |
| | S2 | S |
| | S3 | S |
| leb | S1 | P |
| | S2 | S |
| | S3 | S |
| **Gene mutant** | | |
| $tmc^{1}$ | S1 | P |
| | S2 | S |
| | S3 | S |
| $piezo^{KO}$ | S1 | P |
| | S2 | P |
| | S3 | S |
| $nan^{Gal4}$ | S1 | P |
| | S2 | S |
| | S3 | S |

(Continue to next page)

Continued

| **Gene mutant** | **Substrate** | **Oviposition position** |
|---|---|---|
| *ppk26*$^{Gal4}$ | S1 | P |
| | S2 | S |
| | S3 | P |
| *nompC*$^{3}$ | S1 | P |
| | S2 | S |
| | S3 | S |
| *ppk1*$^{ESB}$ | S1 | P |
| | S2 | S |
| | S3 | S |
| *iav*$^{1}$ | S1 | P |
| | S2 | P |
| | S3 | P |
| *ir8a*$^{1}$ | S1 | P |
| | S2 | P |
| | S3 | P |
